# Exploring the genetic correlations of antisocial behavior and life history traits

**DOI:** 10.1101/247411

**Authors:** Jorim J. Tielbeek, J.C. Barnes, Arne Popma, Tinca J.C. Polderman, James J. Lee, John R.B. Perry, Danielle Posthuma, Brian B. Boutwell

**Author notes:** The authors declare no conflicts of interest. All results reported are original.

## Abstract

Prior evolutionary theory provided reason to suspect that measures of development and reproduction would be correlated with antisocial behaviors in human and non-human species. Behavioral genetics has revealed that most quantitative traits are heritable, suggesting that these phenotypic correlations may share genetic etiologies. We use GWAS data to estimate the genetic correlations between various measures of reproductive development (N= 52,776 – 318,863) and antisocial behavior (N= 31,968). Our genetic correlation analyses demonstrate that alleles associated with higher reproductive output (number of children ever born, r_g_=0.50, p=.0065) were positively correlated with alleles associated with antisocial behavior, whereas alleles associated with more delayed reproductive onset (age of first birth, r_g_=-.64, p=.0008) were negatively associated with alleles linked to antisocial behavior. Ultimately, these findings coalesce with evolutionary theories suggesting that increased antisocial behaviors may partly represent a faster life history approach, which may be significantly calibrated by genes.

---

A moderate proportion of the variance in antisocial phenotypes is accounted for by genetic variation [1]. Phenotypic indicators of externalizing traits, moreover, are correlated with other forms of psychopathological behavior [2]. These findings suggest a general vulnerability underlying the spectrum of externalizing disorders, one underpinned by pleiotropic genetic influences. With the proliferation of population-based samples containing measured genes, researchers have investigated possible molecular correlates of antisocial behaviors. While some promising results emerged, so too did to the recognition of limitations [3–4]. Chief among them included the use of underpowered designs, lack of reproducibility and an absence of correction for multiple testing bias [4].

As a result, studies utilizing genome-wide techniques in large samples have become the preferred approach to unraveling the genetic architectures of complex traits (which have previously been shown to be moderately heritable) [4–10]. Recent GWAS of antisocial phenotypes revealed a number of trait-relevant alleles that were nearly genome-wide significant [5,10]. GWAS evidence has also revealed associations between a number of SNPs and phenotypes known to correlate with antisocial behaviors [6]. In particular, various indicators of reproductive behavior such as age at menarche, age at first sexual contact, and number of sexual partners have all been examined using genetically sensitive data, with results identifying a number of genome-wide significant SNPs [6].

What remains in need of further testing is whether the genetic underpinnings of antisocial behavior covary with the genetic underpinnings of reproductive indicators. For this topic, in particular, prior theoretical work has suggested that some forms of antisocial behavior may represent natural variation in broad reproductive strategies—captured broadly in the area of life history evolution—that have been shaped by natural selection over the course of human evolution [7]. This line of reasoning makes the explicit prediction that reproductive and antisocial behavior should be correlated at a phenotypic level, and perhaps at a genetic level, such that alleles associated with antisocial behavior are also associated with higher reproductive output and more rapid physical maturation. Indeed, in an earlier GWAS study, we examined the genetic covariation of antisocial behavior with a range of cognitive, psychiatric and reproductive outcomes utilizing overlapping data and reported some suggestive genetic correlations with various reproductive traits [5].

We combined GWAS data from the Broad Antisocial Behavior Consortium (BroadABC) with data from the Early Genetics and Life-course Epidemiology (EAGLE) Consortium [10]. In total, we meta-analyzed genotypic and phenotypic data from 31,968 individuals across thirteen unique samples, making it the largest collective sample available to estimate the effects of genome-wide genetic variation on antisocial behavior (ASB). In addition, summary statistics for seven relevant reproductive and longevity traits were obtained from large existing GWAS datasets. The present study explores the genome-wide genetic correlation between reproductive traits and antisocial behavior. By examining the extent to which alleles associated with antisocial behavior are related to alleles underpinning variation in reproductive outcomes, this study could provide a better understanding of why these traits tend to correlate at the phenotypic level [7].

## Method

In the current study, we used (cross-trait) LD-score regression to estimate the SNP-based heritability of ASB and reproductive traits and the genetic correlation between the traits explained by all SNPs [8]. LD-score regression disentangles the contribution of true polygenic signal and bias due to population stratification to the inflated test statistics in GWAS, and optionally calculates a genetic correlation (rg) between traits [8]. An important condition is that the per-SNP heritability near a given SNP must not be confounded with the extent of that SNP’s LD with neighbouring SNPs. This condition is likely to be violated in the case of a phenotype related to fitness producing a downwardly biased SNP-based heritability estimate [8]. Yet, it is still possible for the genetic correlation between two traits to be estimated accurately given the bias in the numerator (the genetic covariance) cancelling the bias in the denominator (the square root of the product of two heritability estimates).

Here, we utilized LD-regression to estimate the genetic correlation of ASB with reproductive traits based on the summary statistics from the largest GWAS meta-analyses available. For reproductive traits, summary statistics were derived from the centralised database LD Hub [9]. Moreover, we employed PLINK to ‘clump’ the SNPs utilizing 1000 Genomes V3 reference panel for Europeans, with 0.1 as LD r2 threshold, and 500 KB as physical distance threshold. Then, we examined whether the signs of the regression coefficients of the SNPs for ASB and age of first birth (yielding the highest r_g_) were, more often than expected by chance, in the same direction. We tested this using a binomial test to verify whether the proportion of SNPs (yielding p-values < .05) with concordant sign was higher or lower than expected by chance (0.5).

For ASB we performed a GWA meta-analysis in order to obtain a large GWAS sample. We combined summary data from the publicly available EAGLE consortium (N= 18,988, [10]) with those from non-overlapping samples of the BroadABC (QIMR, TEDS, COGA, Yale-Penn, N=12,980), totalling 31,968 participants. The genetic correlation between these two meta-analysed datasets was .38 (SE= .48). To maximize sample size, we included studies with a broad range of antisocial measures, including both aggressive and non-aggressive domains of antisocial behaviour, and utilizing study-specific scales in different age groups [5, 10–11]. The meta-analysis was run using a fixed-effects model with z-scores weighted by sample size as implemented in the software METAL [12].

## Results

First, we calculated the SNP-based heritability estimates of ASB and the life-history traits, after which the genetic correlations between the traits were computed. The estimated heritability for ASB utilizing LD-score regression was low—2.8% with a standard error of 1.5%. Substantially higher estimates of the SNP-heritability of ASB have been reported elsewhere using genome-wide complex traits analyses (GCTA) tools. We identified four cohorts (including QIMR, ALSPAC and GENR samples) with ASB measures that previously performed GCTA, resulting in a sample-size weighted *h*^2^ of 35% (Panel A in Table 1). These GCTA estimates are based on more homogeneous individual cohorts yielding smaller samples, which might explain the discrepancy in *h*^2^. The SNP heritabilities based on LD-score regression were 5.9% (SE=2.7%) and 5.2% (SE=2.7%) respectively for the EAGLE and BroadABC consortia. Another possibility is that LD-score regression is more sensitive to causal SNPs being in low LD with their neighbours [12].

**Table 1.** Panel A. Previously reported genome-wide complex trait analysis (GCTA) estimates on ASB.

| Cohort, Trait | N | SNP $H^2$ | SE | P |
| --- | --- | --- | --- | --- |
| *QIMR<sup>1</sup>, adult ASB* | 4816 | .55 | .41 | .07 |
| *ALSPAC<sup>2</sup> (middle childhood/early adolescence)* | 5299 | .08 | .06 | .10 |
| *GENR<sup>2</sup> (6-year)* | 2101 | .54 | .19 | .002 |
| *NTR<sup>2</sup> (3-year)* | 908 | .46 | .35 | .09 |

| *Phenotype* | *Sample Size* | *SNP $h^2$* | *$r_g$(SE)* | *Bivariate Intercept* | *P-value* |
| --- | --- | --- | --- | --- | --- |
| *Age at Menarche* | 182.416 | .21 | .026(.085) | -0,0083 | .76 |
| *Age of First Birth* | 222.037 | .063 | -.64(.19) | -0,0003 | .0008** |
| *Number of Children Ever Born* | 318.863 | .025 | .50(.18) | -0,007 | .0065** |
| *Age at Menopause* | 69.360 | .13 | -.10(.144) | -0,0016 | .48 |
| *Mother's Age at Death* | 52.776 | .040 | -.51(.26) | -0,0017 | .045* |
| *Father's Age at Death* | 63.775 | .042 | -.52(.24) | -0,0036 | .026* |
| *Parents' Age at Death* | 45.627 | .032 | -.70(.29) | 0,0093 | .015* |

Note: \*Nominally significant ( $p < .05$ ) \*\*Significant at the multiple-testing corrected p-value (.0071). GWAS summary statistics from our combined analyses were used to calculate the $r_g$ 's with other traits. SNP $h^2$ is the estimation of narrow-sense heritability contributed by common SNPs. $r_g$ is the genetic correlation and is calculated with the LDSC software package using pre-calculated LD scores from Bulik-Sullivan et al.

Our main analysis (Panel B in Table 1, Figure 1) revealed significant (α<0.0071) genetic correlations of ASB with age of first birth (r_g_=-.64, p=.0008) and number of children ever born (r_g_=0.50, p=.0065) as well as suggestive associations with parents’, mothers’ and fathers’ age at death, but not with age at menarche or age at menopause (r_g_=.026, p=.76; r_g_=-.10, p=.48, respectively). Moreover, sign-tests revealed less SNPs yielding a p-value < .05 with the same direction of effect than expected by chance (proportion = .47, p <.001) for ASB and age of first birth.

**Figure 1.**
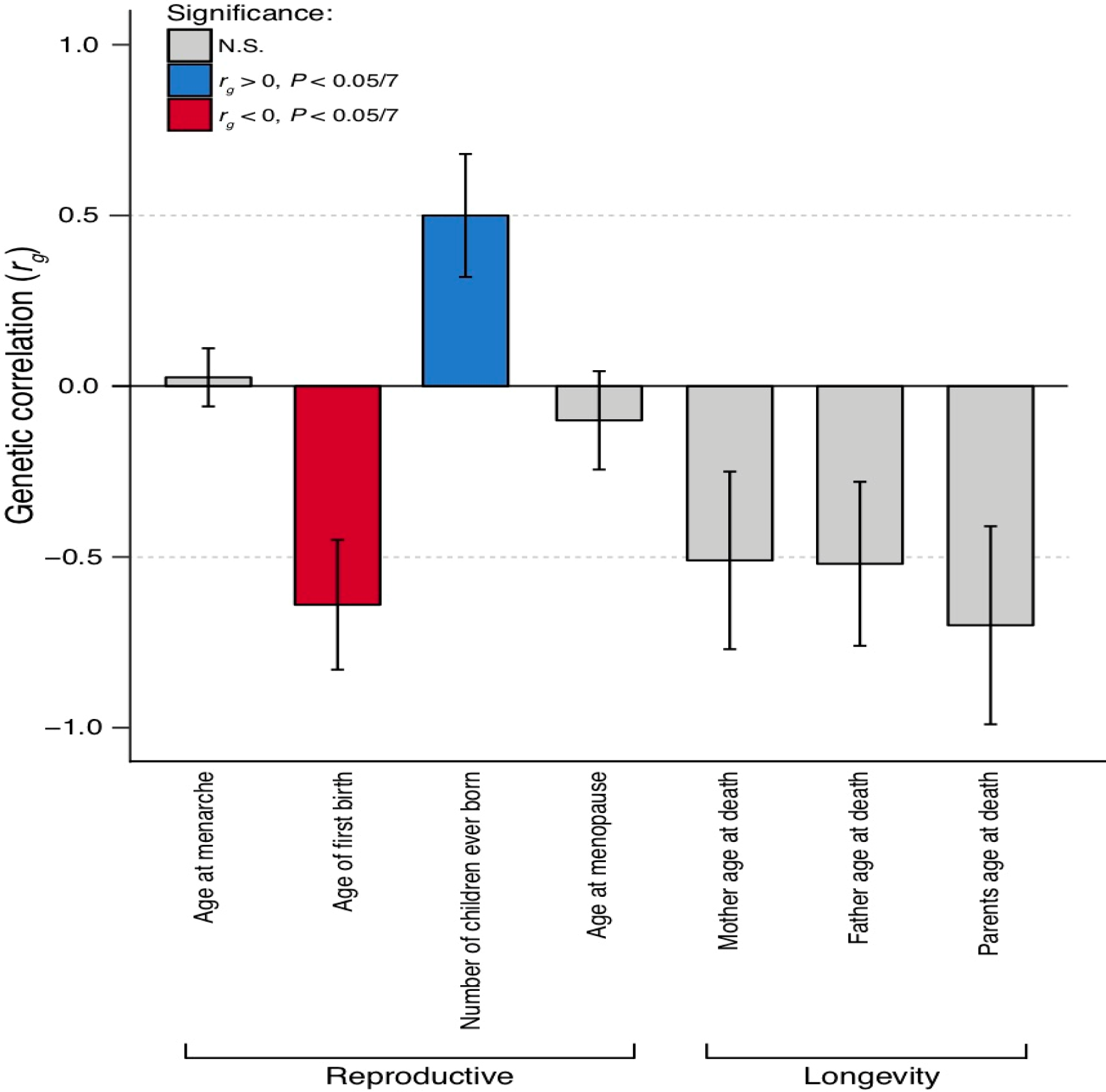
Genetic Correlations between Antisocial Behavior and Life History Variables.

## Discussion

The present study expands on prior work [5], showing preliminary evidence of a genetic correlation between reproductively relevant traits and antisocial behavior. Our genetic correlation analyses demonstrate that alleles associated with higher reproductive output (i.e., faster life history styles) were positively correlated with alleles associated with antisocial behavior, whereas alleles associated with giving birth later in life were negatively associated with alleles linked to antisocial behavior. It is important to acknowledge that this study is preliminary and not without limitations. Our correlations were somewhat higher than expected (ranging from 0.5 to 0.7). We hypothesize that with increasing samples and more homogeneous phenotypic measures these estimates will be more in the range of 0.3-0.5, as reported by previous studies [5].

Our genetic correlation analyses are based on, and limited by, relatively low SNP-based heritability estimates of ASB (2.9%), yielding a heritability Z score of 1.93. The low SNP-based *h*^2^ estimate of ASB may be due to heterogeneity in measurement by meta-analyzing multiple cohorts or the tendency of the causal SNPs to be in low LD with their neighbors. Finally, although somewhat elevated, the magnitude of the genetic correlations observed herein is similar in magnitude compared to other related phenotypes, utilizing alternative samples [5,14]. These results, although promising and guided by prior theory [7], should be viewed only as preliminary. As sample sizes continue to grow, and a more diverse range of phenotypes become available for testing, additional insight should emerge regarding the molecular underpinnings for antisocial behavior and human reproductive strategy.

## Acknowledgements

The authors would like to thank Philip R Jansen for technical assistance.

## References

1. Polderman TJ, Benyamin B, De Leeuw CA, Sullivan PF, Van Bochoven A, Visscher PM, Posthuma D. Meta-analysis of the heritability of human traits based on fifty years of twin studies. Nat Genet 2015; 47: 702–709.

2. Krueger RF, Hicks BM, Patrick CJ, Carlson SR, Iacono WG, McGue M. Etiologic connections among substance dependence, antisocial behavior and personality: Modeling the externalizing spectrum. J Abnorm Psychol 2002; 111: 411.

3. Dick DM, Agrawal A, Keller MC, Adkins A, Aliev F, Monroe S, Sher KJ. Candidate gene–environment interaction research: Reflections and recommendations. Perspec Psychol Sci 2015; 10: 37–59.

4. Chabris CF, Lee JJ, Cesarini D, Benjamin DJ, Laibson DI. The fourth law of behavior genetics. Curr Dir Psychol Sci 2015; 24: 304–312.

5. Tielbeek JJ, Johansson A, Polderman TJ, Rautiainen MR, Jansen P, Taylor M, Viding E. Genome-Wide Association Studies of a Broad Spectrum of Antisocial Behavior. JAMA Psychiatry 2017.

6. Barban N, Jansen R, De Vlaming R, Vaez A, Mandemakers JJ, Tropf FC, Tragante V. Genome-wide analysis identifies 12 loci influencing human reproductive behavior. Nat Genet 2016; 48: 1462–1472.

7. Boutwell BB, Barnes JC, Beaver KM, Haynes RD, Nedelec JL, Gibson CL. A unified crime theory: the evolutionary taxonomy. Aggress Violent Behav 2015; 25: 343–353.

8. Bulik-Sullivan BK, Loh PR, Finucane HK, Ripke S, Yang J, Patterson N, Schizophrenia Working Group of the Psychiatric Genomics Consortium. LD Score regression distinguishes confounding from polygenicity in genome-wide association studies. Nat Genet 2015; 47: 291–295.

9. Zheng, et al. LD Hub: a centralized database and web interface to perform LD score regression that maximizes the potential of summary level GWAS data for SNP heritability and genetic correlation analysis. Bioinformatics 2017 5; 272–279.

10. Pappa I, St Pourcain B, Benke K, Cavadino A, Hakulinen C, Nivard MG, Evans DM. A genome-wide approach to children’s aggressive behavior: the EAGLE consortium. Am J Med Genet B: Neuropsychiatr Genet 2016; 171: 562–572.

11. Tielbeek JJ, Medland SE, Benyamin B, Byrne EM, Heath AC, Madden PA, Verweij KJ. Unraveling the genetic etiology of adult antisocial behavior: a genome-wide association study. PloS one 2012; 7: e45086.

12. Willer CJ, Li Y, Abecasis GR. METAL: fast and efficient meta-analysis of genomewide association scans. Bioinformatics 2010; 26: 2190–2191.

13. Lee JJ, Chow CC. Conditions for the validity of SNP-based heritability estimation. Hum Genet 2014; 133: 1011–1022.

14. Hill WD, Marioni RE, Maghzian O, Ritchie SJ, Hagenaars SP, McIntosh AM, Deary IJ. A combined analysis of genetically correlated traits identifies 187 loci and a role for neurogenesis and myelination in intelligence. Molec Psychiat 2018; 1.

